# Structural basis for Nirmatrelvir in vitro efficacy against SARS-CoV-2 variants

**DOI:** 10.1101/2022.01.17.476556

**Authors:** Samantha E Greasley, Stephen Noell, Olga Plotnikova, Rose Ann Ferre, Wei Liu, Ben Bolanos, Kimberly Fennell, Jennifer Nicki, Tim Craig, Yuao Zhu, Al E Stewart, Claire M Steppan

## Abstract

The COVID-19 pandemic continues to be a public health threat with emerging variants of SARS-CoV-2. Nirmatrelvir (PF-07321332) is a reversible, covalent inhibitor targeting the main protease (Mpro) of SARS-CoV-2 and the active protease inhibitor in PAXLOVID^™^ (nirmatrelvir tablets and ritonavir tablets). We evaluated the *in vitro* catalytic activity and *in vitro* potency of nirmatrelvir against the main protease (M^pro^) of prevalent variants of concern (VOC) or variants of interest (VOI): Alpha (α, B.1.1.7), Beta (β, B.1.351), Delta (δ, B1.617.2), Gamma (*γ*, P.1), Lambda (λ, B.1.1.1.37/C37), Omicron (o, B.1.1.529) as well as the original Washington or wildtype strain. These VOC/VOI carry prevalent mutations at varying frequencies in the M^pro^ specifically for: α, β, *γ* (K90R), λ (G15S) and o (P132H). In vitro biochemical enzymatic assay characterization of the enzyme kinetics of the mutant M^pros^ demonstrate that they are catalytically comparable to wildtype. Nirmatrelvir has similar potency against each mutant M^pro^ including P132H that is observed in the Omicron variant with a Ki of 0.635 nM as compared to a Ki of 0.933nM for wildtype. The molecular basis for these observations were provided by solution-phase structural dynamics and structural determination of nirmatrelvir bound to the o, λ and β Mpro at 1.63 - 2.09 Å resolution. These in vitro data suggest that PAXLOVID has the potential to maintain plasma concentrations of nirmatrelvir many-fold times higher than the amount required to stop the SARS-CoV-2 VOC/VOI, including Omicron, from replicating in cells (1).

## Introduction

New viral infectious diseases are emerging and have caused major public health crises in recent years. Reported examples in the last 20 years include severe acute respiratory syndrome coronavirus (SARS-CoV), H1N1 influenza, the Middle East respiratory syndrome coronavirus (MERS-CoV), Ebola virus disease (EVD), and Zika virus (2). The world continues to grapple with a global pandemic caused by a novel coronavirus, SARS-CoV-2 that was initially reported to the World Health Organization (WHO) on December 31, 2019 (3). Later, WHO designated this virus as the severe acute respiratory syndrome coronavirus-2 (SARS-CoV-2) owing to its similarity with the previous SARS-CoV (4). Then at the end of January 2020, WHO declared this viral outbreak as a public health emergency of international concern (5), and subsequently characterized it as a pandemic.

Genetic lineages of SARS-CoV-2 have been emerging and circulating around the world since the beginning of the COVID-19 pandemic. Like all viruses, SARS-CoV-2 is constantly changing through mutation and each virus with a unique sequence is considered a new variant. The World Health Organization as well as other public health organizations monitor all variants that cause COVID-19 for increased risk to global public health and classify as variants being monitored, variants of interest (VOI), variants of concern (VOC) and variants of high consequence. As of February 3, 2022, the WHO has designated five VOCs: B.1.1.7 (Alpha, α), B.1.351 (Beta, β), P.1 (Gamma, *γ*), B.1.617.2 (Delta, δ), and B.1.1.529 (Omicron, o) while B.1.1.1.37 (Lambda, λ) and B.1.621 (Mu, μ) have been designated as ‘variants of interest,’ VOI (6).

SARS-CoV-2 is a highly infectious beta coronavirus that can be life-threatening in serious cases. While effective COVID-19 vaccines have been developed, for individuals who have yet to be vaccinated or cannot be vaccinated, such as due to pre-existing medical conditions, therapeutics will likely be needed to effectively combat coronavirus disease 2019 (COVID-19). (1)

SARS-CoV-2 main protease (M^pro^, also referred to as 3CL protease) is a cysteine protease that is critical for the processing of the two polyproteins (pp1a and pp1ab) encoded by the SARS-CoV-2 genome. The protease cleaves these polyproteins into shorter, non-structural proteins that are essential for viral replication (1). Owing to this key role in viral replication, small molecule inhibitors of SARS-CoV-2 M^pro^ represent attractive therapeutics for the treatment of COVID-19. We have previously reported on the discovery and antiviral efficacy of nirmatrelvir (PF-07321332), an orally bioavailable SARS-CoV-2-M^pro^ inhibitor with in vitro pan-human coronavirus antiviral activity with excellent off-target selectivity and in vivo safety profiles (1). Nirmatrelvir has demonstrated oral activity in a mouse-adapted SARS-CoV-2 model and has achieved oral plasma concentrations that exceed the in vitro antiviral cell potency, in a phase I clinical trial in healthy subjects (1). Here we report on the catalytic activity of frequently observed M^pro^ mutations in SARS-CoV-2 VOC/V OI, in vitro efficacy of nirmatrelvir against these mutant M^pros^, the solution-phase structural dynamics and structure of nirmatrelvir bound to the M^pros^ from three VOCs, β, λ and o.

## Results

Full-length wildtype M^pro^ from the original Washington variant (USA-WA1/2020) and VOC/VOI SARS-CoV-2 M^pro^ were expressed and purified to near homogeneity as demonstrated by a singular protein peak with a confirmed intact mass of 33.8 kDa for a fully authentic form of each protein (Figure 1A). A final and size exclusion chromatography step showed the wildtype, K90R, G15S and P132H M^pro^ proteins to be nearly 100% pure by Western blot analysis (Figure 1B). An established M^pro^ fluorescence resonance energy transfer (FRET)- based cleavage assay was used to determine enzyme catalytic activity by monitoring initial velocities of the proteolytic activities at varying substrate (SARS canonical peptide) concentrations (1,7,8) (Figure 1C). The turnover number (kcat) and Michaelis constants (Km) was determined for the wildtype, K90R, G15S and P132H M^pro^ proteins, respectively ((Table 1). The catalytic efficiencies (kcat/Km) of the K90R (28255 S^-1^M^-1^), G15S (16483 S^-1^M^-1^), and P132H (22692 S^-1^M^-1^) are similar to wildtype Mp^ro^ (39830 S^-1^M^-1^). These data suggest that the K90R, G15S and P132H M^pro^ variants exhibit comparable enzymatic activities as compared to wildtype M^pro^. Next, we evaluated the ability of nirmatrelvir to inhibit wildtype and K90R, G15S and P132H M^pro^ enzymatic activities (Figure 1D). Nirmatrelvir potently inhibited wildtype (mean Ki of 0.93 nM) and the mutated enzymes containing the K90R (Ki 1.05nM), G15S (Ki 4.07 nM) and P132H (Ki 0.64 nM) M^pro^ (Table 2). A comparison of the VOC/VOI enzyme potency to that of wildtype M^pro^ by t-Test using the log of individual Ki values with one tail and un-equal variance determined that the Ki values are not statistically different for the K90R and P132H M^pro^ proteins. There was a statistically significant shift in potency for G15S Mpro mutant, frequently observed in the Lambda variant relative to wildtype (P<0.0005).

**Figure 1.**
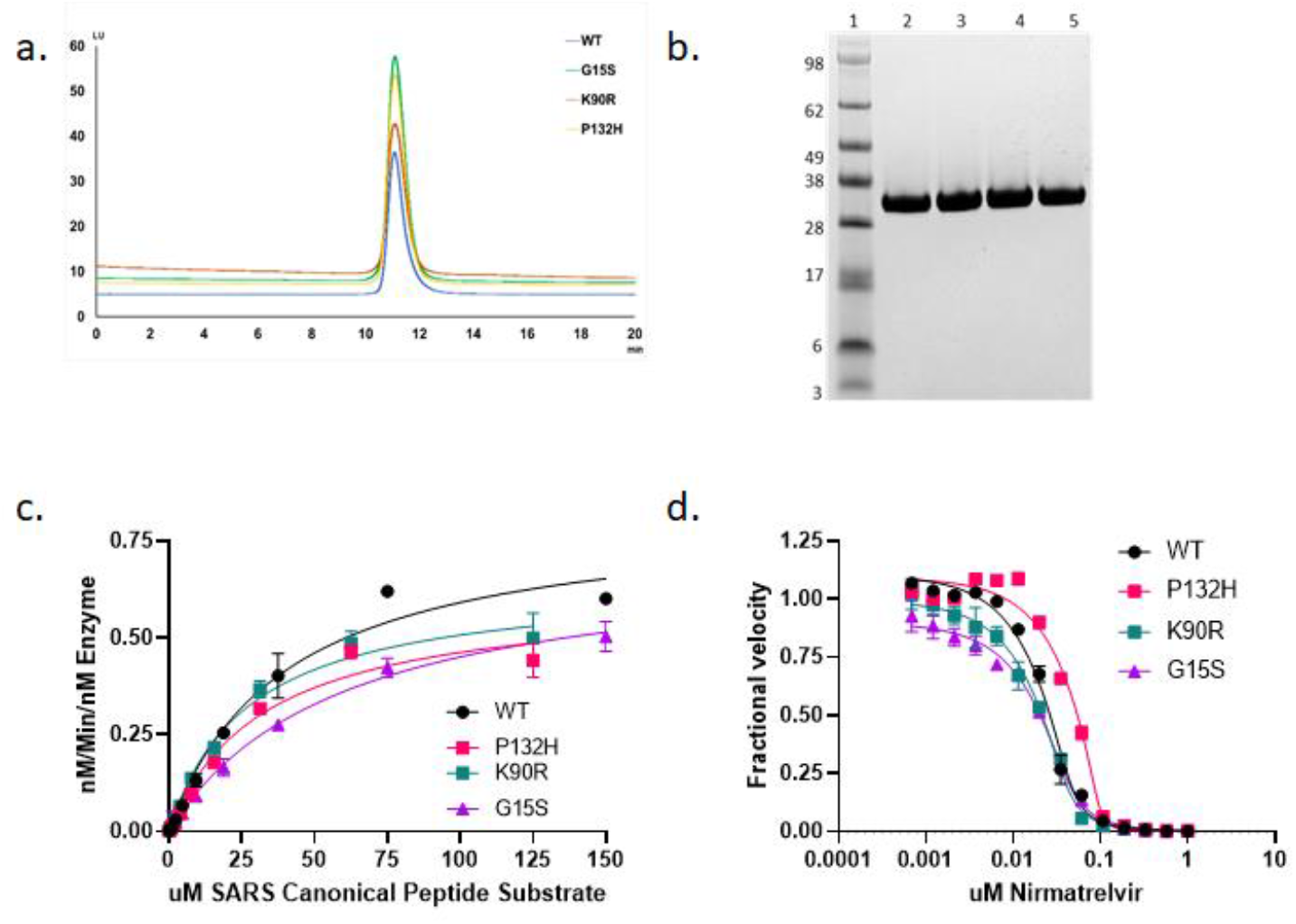
Purification and characterization of SARS-Cov2-M^pro^ P132H. (a) Analytical size-exclusion profile using using Zenix SEC-300, 4.6 x 150 mM column (b) SDS-PAGE analysis: Lane 1, Marker; Lane 2, SARS-Cov2-M^pro^ wildtype; Lane 3, SARS-Cov2-M^pro^ G15S; Lane 4, SARS-Cov2-M^pro^ K90R; Lane 5, SARS-Cov2-M^pro^ P132H (c) Enzyme Kinetics of M^pro^ variants: The rate of cleavage of the FRET peptide substrate in the presence of indicated M^pro^ is monitored by increase in fluorescence over time with the fluorescent signal being converted to nM substrate cleaved by use of a standard curve generated from cleaved substrate. The data was then normalized to the amount of enzyme used in the experiment. (d) Enzyme Inhibition of M^pro^: M^pro^ activity is monitored in the presence of increasing concentrations of nirmatrelvir with Ki values calculated using the Morrison equation to fit the data.

**Table 1:**
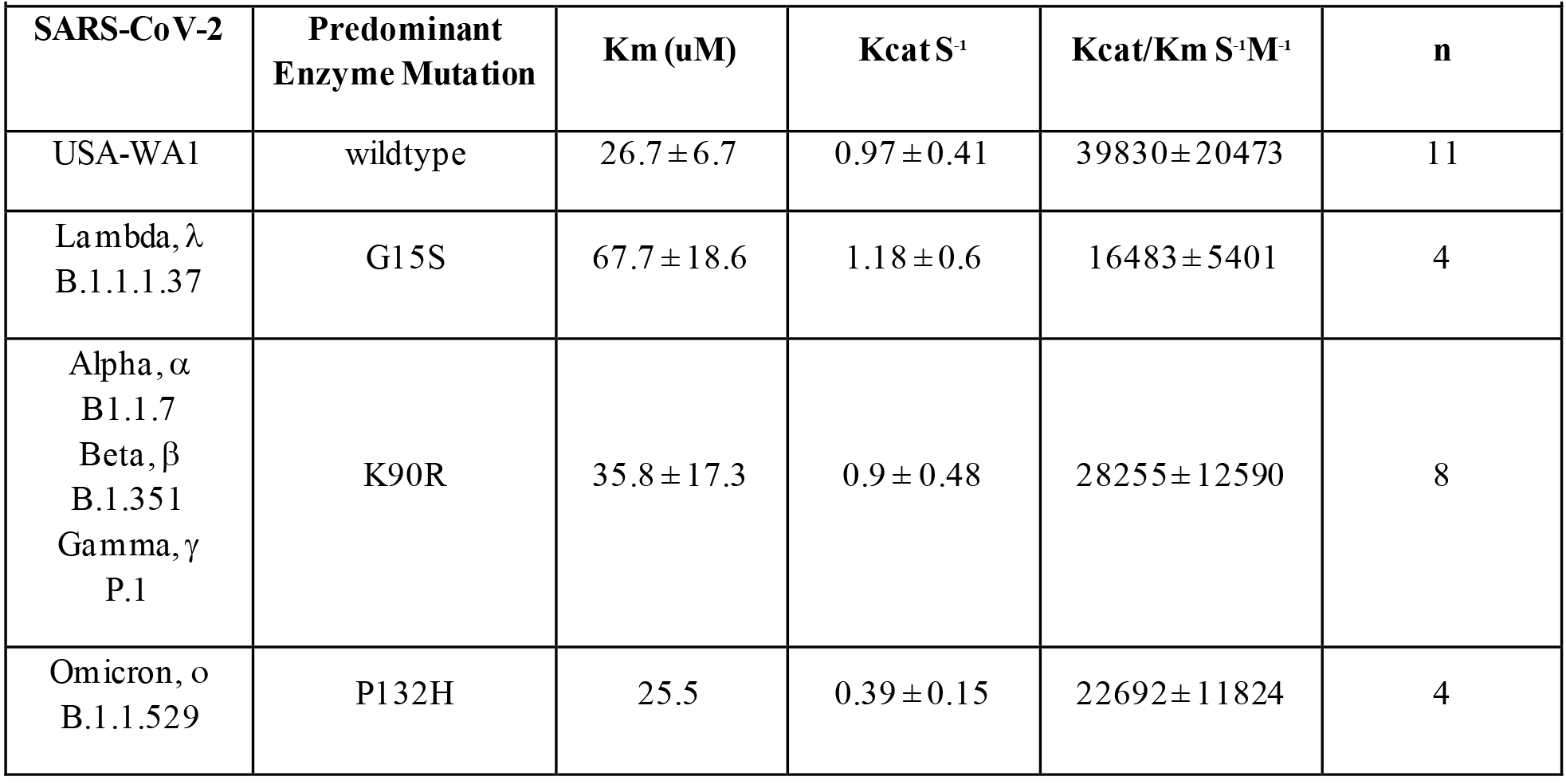
Enzymatic Catalytic Constants for SARS-CoV2 M^pro^ Enzyme.

| Enzymatic Catalytic Constants for SARS-CoV2 M<sup>pro</sup> Enzyme. |  |  |  |  |  |
| --- | --- | --- | --- | --- | --- |
| SARS-CoV-2 | Predominant Enzyme Mutation | K<sub>m</sub> (uM) | K<sub>cat</sub> S<sup>-1</sup> | K<sub>cat</sub>/K<sub>m</sub> S<sup>-1</sup>M<sup>-1</sup> | n |
| USA-WA1 | wildtype | 26.7 ± 6.7 | 0.97 ± 0.41 | 39830 ± 20473 | 11 |
| Lambda, λ<br>B.1.1.1.37 | G15S | 67.7 ± 18.6 | 1.18 ± 0.6 | 16483 ± 5401 | 4 |
| Alpha, α<br>B1.1.7<br>Beta, β<br>B.1.351<br>Gamma, γ<br>P.1 | K90R | 35.8 ± 17.3 | 0.9 ± 0.48 | 28255 ± 12590 | 8 |
| Omicron, o<br>B.1.1.529 | P132H | 25.5 | 0.39 ± 0.15 | 22692 ± 11824 | 4 |

**Table 2:** In Vitro Potency of Nirmatrelvir for Inhibiting the SARS-CoV2 Mutant M^pro^ Activity in a FRET Assay.

|  |  |  |  |  |  |
| --- | --- | --- | --- | --- | --- |
| **Mutant Enzyme** | **Ki (nM) GeoMean** | **Ki (nM) lower 95% CI** | **Ki (nM) upper 95%** | **n** | **P-value<sup>b</sup> to Wild Type Ki** |
| G15S | 4.07 | 2.62 | 6.32 | 4 | 0.0002 |
| K90R | 1.05 | 0.19 | 5.82 | 5 | 0.371 |
| P132H | 0.635 | 0.179 | 2.25 | 4<sup>a</sup> | 0.074 |
| wild type | 0.933 | 0.471 | 1.85 | 9<sup>a</sup> | NA |

a. The n value represents the number of Ki values used to determine the geomean and CI which is lower than the experiment count due to censoring, ie experimental values that are < are excluded from GeoMean calculation.
b. p-value calculated as a t-test statistic for log Ki values compared to wild type.

The crystal structures of nirmatrelvir bound to the three VOC were determined to 2.09 Å (K90R), 1.68 Å (G15S) and 1.63 Å (P132H) resolution (Figure 2). Superposition of each of these mutant structures with the wildtype M^pro^, the structure of which was reported in detail previously (5,14), shows that the binding mode of nirmatrelvir is unperturbed by the mutations, with the ligand maintaining the protein interactions observed in the wildtype M^pro^, as shown in Figure 2. Indeed, these mutations are distal to the PF-07321332 binding pocket, with Pro 132 located approximately 16 Å (Cα-Pro 132 to Cα-Glu166) from the binding pocket, while K90R and G15S reside 19 Å and 17 Å respectively (Cα-R90/S15 to Cα-Cys145) from the PF-07321332 binding pocket (Figure 2D). The crystal structures also show that the mutations do not give rise to any signification changes of the protein around the binding pocket or the site of the mutation (Figure 2D)

**Figure 2.**
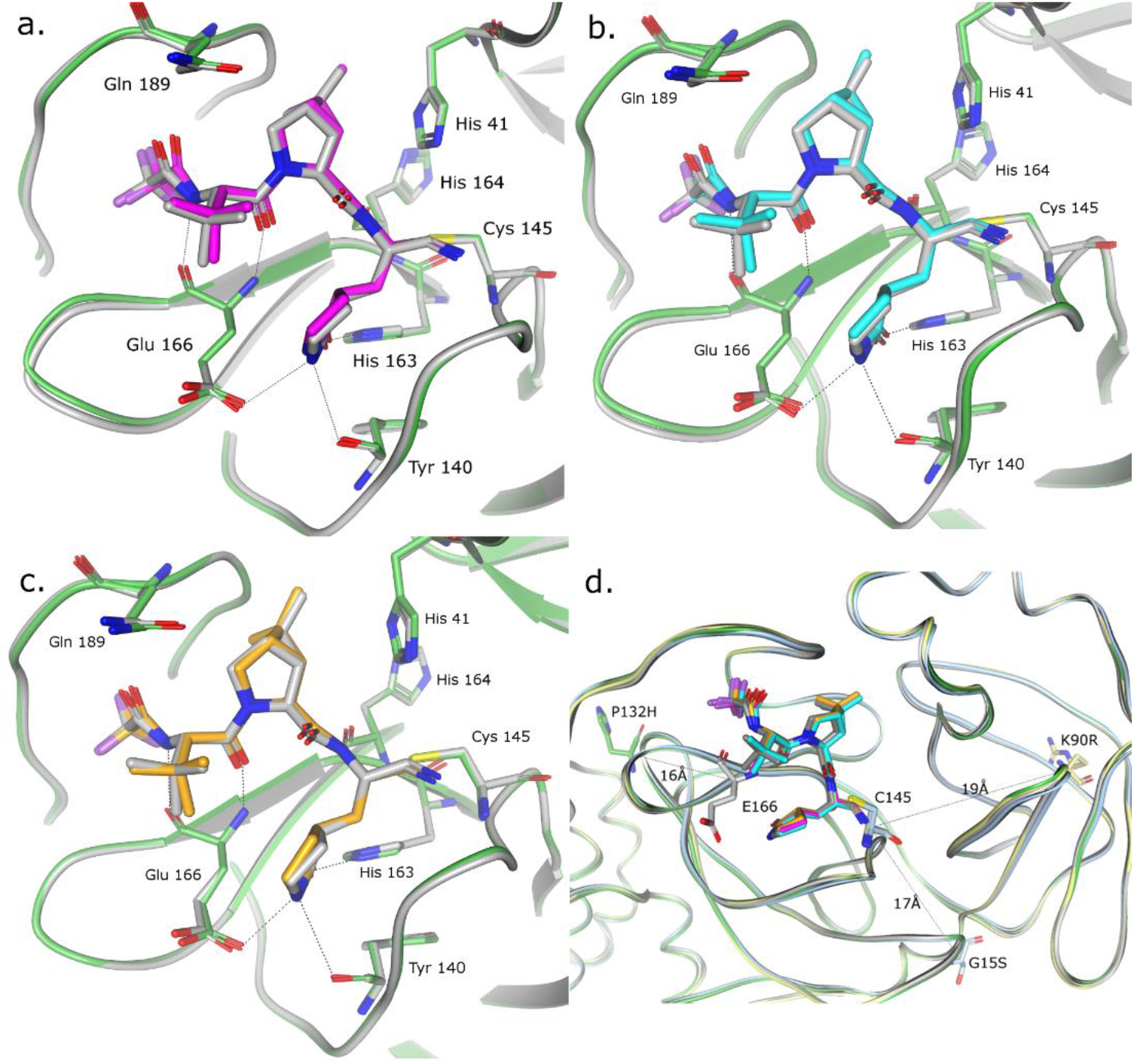
Structural Characterization of nirmatrelvir bound to SARS-Cov-2-M^pro^ VOC. Superposition of the x-ray crystal structures of nirmatrelvir bound to wildtype SARS-CoV-2-M^pro^ (grey) and (a.) SARS-Cov-2-M^pro^ P132H (in magenta and green); (b.) G15S (in cyan and green) and (c.) K90R (in orange and green). (d.) Ribbon figure of the 3 VOC crystal structures superposed on the WT, highlighting the location and distance of the mutant residues (in stick representation) in relation for the nirmatrelvir binding site. Colors are maintained from figures 2 a,b and c.

To provide an extended SARS-Cov-2 M^pro^ structural assessment, the solution-phase structural dynamics of K90R, G15S and P132H along with the wildtype SARS-Cov2 M^pro^ (10 μM, 25 mM Tris pH=7.2, 150 mM NaCl) were individually profiled using HDX-MS. A residual plot (Figure 3) comparing wildtype and mutant deuterium uptake profiles (HD Examiner 3.3.0 software, Sierra Analytics) revealed no significance differences (+ 6% deuterium) in the backbone dynamics of K90R, G15S or P132H from wild-type SARS-Cov-2 M^pro^. These results also suggest that the mutations alone did not produce a significant shift from wildtype solution-phase backbone conformational dynamics.

**Figure 3.**
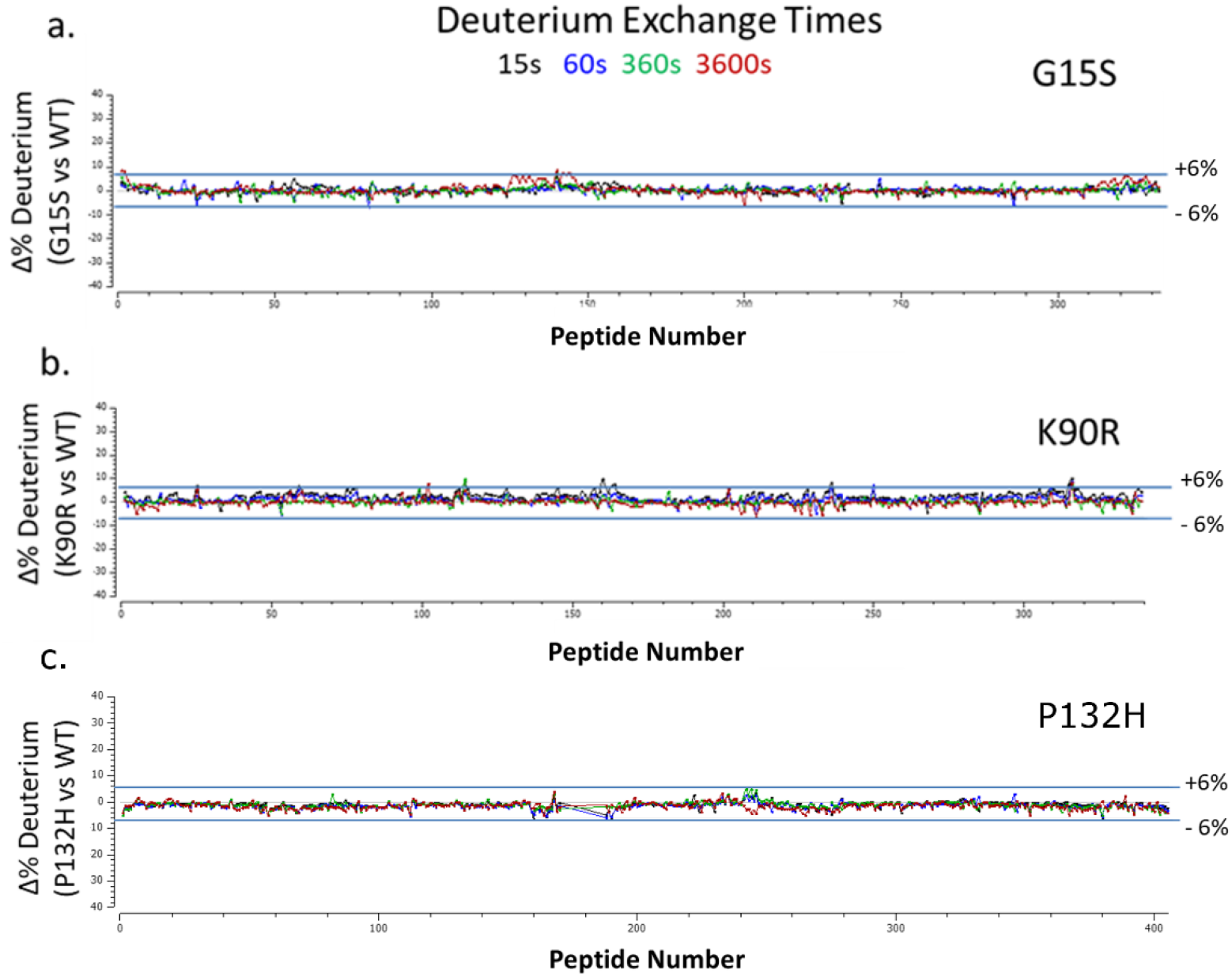
Residual deuterium exchange plots. indicate no significant differential uptake between wild-type and SARS-CoV-2-M^pro^ mutants; (a.) G15S, (b.) K90R and (c.) P132H.

## Discussion

We evaluated the catalytic activity and potency of nirmatrelvir against the Mpro mutations observed in the prevalent variants of concern (VOC): Alpha (α, B.1.1.7), Beta (β, B.1.351), and Gamma (*γ*, P.1), which harbor a K90R mutation; Lambda (λ, B.1.1.1.37/C.37) with a G15S mutation and Omicron (o, B.l.1.529) with a P132H mutation. The K90R, G15S and P132H M^pro^ mutations exhibited comparable enzymatic properties as compared to wildtype M^pro^ as evidenced by our determined catalytic efficiencies (kcat/Km). We demonstrate that nirmatrelvir has a comparable potency against these mutant M^pros^ relative to wildtype (M^pro^ from the original Washington strain (USA-WA1/2020)). Furthermore, deuterium exchange profiles of K90R, G15S and P132H showed equivalent uptake to wildtype M^pro^, and revealed no detectable differences in backbone dynamics. In combination with static high-resolution crystal structures, HDX-MS exchange from the ensemble of solution-phase conformations, provides evidence of equivalent solution-phase backbone dynamics for K90R, G15S and P132H to wildtype and suggests no evidence for influence from crystal lattice packing in the crystal structures Indeed, the x-ray crystal structures of nirmatrelvir bound to K90R, G15S and P132H M^pro^ variants shows that, not only is the binding mode of nirmatrelvir indistinguishable to the wildtype M^pro^ structure, but also that the conformation of the protein is not altered by the mutations, in agreement with the solution-phase data. These observations are consistent with recently reported biochemical characterization with similar kcat/Km and nirmatrelvir potency values (10, 11). Nirmatrelvir also appears to retain its in vitro antiviral efficacy against Omicron relative to wildtype (12–15).

The emergence of naturally occurring SARS-CoV-2 variants exemplify its ability to mutate and signify the continued potential for this pandemic to be problematic. It is important to continue monitoring emerging variants of concern to quickly understand potential challenges to theefficacy of current and future anti-viral therapies. More broadly, the availability of naturally occurring mutations could provide opportunity to more deeply understand additional aspects of protease structure-function and fitness. Studies described here demonstrate the in vitro inhibitory activity of nirmatrelvir against the Alpha, Beta, Gamma, Lambda and Omicron variants of M^pro^ and indicate the structural basis for retention of in vitro potency against these mutant proteins. They also inform the methods for assessing activity against subsequent variants possessing mutations in the M^pro^ protein.

## Experimental procedures

### Protein production and purification

Briefly, the wild type SARS-Cov2-M^pro^ construct was designed based on Su et al 2020 (16). An additional N-terminal PreScission protease cleavage site was inserted between GST and the self - cleavage site. Site directed mutagenesis was performed to make each of the variants (K90R, G15S or P132H). The resulting plasmid was then transformed into BL21 (DE3) cells for protein expression. One liter of LB media was inoculated with 30mL of overnight culture and grown at 37°C until an OD_600_ of 0.6 was reached. The culture was induced using a final concentration of 0.2 mMIPTG and harvested 1 hour post induction. Cells were lysed in 20 mM Tris pH 8.0 buffer containing 500 mM NaCl, 10% glycerol, 0.2 mM TCEP with a microfluidizer, and the mixture was clarified by centrifugation at 15000 x g. The resulting supernatant was purified by a Ni-affinity column using a step gradient, followed by C-terminal His-tag cleavage with PreScission protease, and a secondary Ni-affinity purification to remove non-cleaved M^pro^ and PreScission protease.

### Enzyme characterization and kinetics

The enzymatic activity of the main protease, M^pro^ of SARS CoV-2 wildtype and variants was monitored using a continuous fluorescence resonance energy-transfer (FRET) assay (1,7,8). The SARS CoV-2 M^pro^ assay measures the activity of full-length SARS CoV-2 M^pro^ protease to cleave a synthetic fluorogenic substrate peptide with the following sequence DABCYL-KTSAVLQ-SGFRKME-EDANS modeled on a consensus peptide. The fluorescence of the cleaved EDANS peptide (excitation 340 nm/emission 490 nm) is measured using a fluorescence intensity protocol on a PHARAstar FSX^™^ microplate reader (BMG Labtech). The assay reaction buffer contained 20 mM Tris–HCl (pH 7.3), 100 nM NaCl, 1 mM EDTA, 5 mM TCEP. 30 nM SARS CoV-2 M^pro^ mutant protease was added to low volume 384-well plates and enzyme reactions were initiated with the addition of 30 μM peptide substrate. Enzyme kinetic constants were calculated from initial rates with the RFU/sec slope converted to uM/sec activity from a standard curve of cleaved substrate. Enzyme inhibition was calculated from the RFU/sec slope collected over 15 min and expressed as percent activity based on control wells containing no compound representing 100% activity and wells containing no enzyme representing 0% activity. Compounds were spotted into assay plates using an Echo 555 acoustic liquid handler in a direct dilution scheme that results in a 1 uM top dose with 1.75-fold dilutions over 14 points. Ki values were fit to the Morrison equation with the enzyme concentration parameter fixed to 30 nM, the *K*_m_ parameter fixed to the substrate Km determined for each mutant and the substrate concentration parameter fixed to 30 μM.

### Crystallization and structure determination

Apo crystals of SARS-CoV-2 M^pro^ mutants were obtained via vapor diffusion in sitting drops using MRC-Maxi (Swissci) plates where protein, at 7.30 mg/ml, was mixed 1:1 with well solution containing 20-24% w/v polyethylene glycol (PEG) 3350 and 0.12-0.21 M sodium sulfate. Plates were incubated at 21°C and crystals grew in under 24 hours. PF-07321332 (1 mM final concentration) was added directly to the crystallization drops and allowed to soak into the apo crystals for 3 hours at 30°C. Soaked crystals were then passed through a cryoprotectant consisting of well buffer containing 20% ethylene glycol, and flash cooled in liquid nitrogen in preparation for data collection.

X-ray diffraction data were collected at −173°C at IMCA-CAT 17-ID beamline (17, 18)of the Advanced Photon Source (APS) at Argonne National Labs (19) using the Eiger 2 x 9M detector (Dectris). The structures were determined by difference Fourier and refined using the anisotropically scaled data as described previously for wildtype SARS-CoV-2-M^pro^ in complex with PF-07321332 (1). Diffraction data processing and model refinement statistics for each of the mutant M^pro^ are given in Table 3.

**Table 3:** Data Statistics for X-ray diffraction data.

| **Data Statistics for X-ray diffraction data** |  |  |  |
| --- | --- | --- | --- |
| **Mutant** | **P132H** | **G15S** | **K90R** |
| PDB entry ID | 7TLL | xxxx | xxxx |
| Wavelength (Å) | 1.0 | 1.0 | 1.0 |
| Resolution | 112.84 – 1.63 | 112.33-1.68 | 110.22-2.09 |
| Space group | P2<sub>1</sub> | P2<sub>1</sub> | P2<sub>1</sub> |
| Unit cell dimensions [Å] | a = 45.4, b = 53.8, c = 115.5 | a = 45.6, b = 53.8, c = 114.3 | a = 45.5, b = 55.8, c = 114.2 |
| Unit cell dimensions [°] | $\alpha = \gamma = 90.0$, $\beta = 102.4$ | $\alpha = \gamma = 90.0$, $\beta = 100.8$ | $\alpha = \gamma = 90.0$, $\beta = 105.2$ |
| Total number of reflections<sup>a</sup> | 178785 (6869) | 156159 (6075) | 95920 (4735) |
| Unique reflections<sup>a</sup> | 53153(2659) | 46261 (2314) | 28383 (1419) |
| Multiplicity<sup>a</sup> | 3.4 (2.6) | 3.4 (2.6) | 3.4 (3.3) |
| Completeness (%), spherical<sup>a</sup> | 78.0 (18.9) | 74.2 (23.9) | 85.7 (35.2) |
| Completeness (%), ellipsoidal<sup>a</sup> | 92.8 (53.2) | 81.0 (46.4) | 87.4 (40.6) |
| Mean I/$\sigma$(I)<sup>a</sup> | 11.5 (1.4) | 10.5 (1.5) | 7.9 (1.6) |
| $R_{\text{merge}}^b$ | 0.056 (0.607) | 0.064 (0.595) | 0.117 (0.789) |
| $R_{\text{pim}}^c$ | 0.035 (0.453) | 0.041 (0.454) | 0.075 (0.499) |
| $CC_{1/2}^d$ | 0.999 (0.635) | 0.998 (0.601) | 0.996 (0.583) |
| **Refinement Statistics** |  |  |  |
| Reflections used | 53153 | 46261 | 28383 |
| Reflections used for $R_{\text{free}}$ | 2620 | 2222 | 1502 |
| $R_{\text{cryst}}^e$ | 0.211 | 0.208 | 0.204 |
| $R_{\text{free}}$ | 0.250 | 0.248 | 0.266 |
| **Ramachandran Plot** |  |  |  |
| Favoured regions (%) | 98.2 | 98.0 | 97.5 |
| Allowed Regions (%) | 1.5 | 1.5 | 2.2 |
| Outlier regions (%) | 0.3 | 0.5 | 0.3 |

<sup>a</sup> Numbers in parentheses refer to the highest resolution shell
$R_{\text{free}}$ is the same as $R_{\text{cryst}}$ , but for 5% of the data randomly omitted from refinement. (22)

### Hydrogen Deuterium exchange

Native protein was deuterium exchanged at four time points (15s, 1m, 6m, 1h), and subsequently digested with two protease columns (Protease XIII/Pepsin (NovaBioAssays) and Nepenthesin-1 (AffiPro) to generate deuterated peptides for LC-MS analysis. A total of 405 WT peptides were identified (Table 4) on the Fusion Lumos OrbiTrap mass spectrometer (mass tolerance <5 ppm) using Proteome Discoverer 2.2.0 software (Thermo Fisher Scientific).

**Table 4.** SARS-CoV-2 Deuterium Exchange Summary (Wild type, P132H, G15S, K90R M^pro^)

| **SARS-CoV-2 Deuterium Exchange Summary (Wild type, P132H, G15S, K90R M<sup>pro</sup>)** |  |  |  |  |
| --- | --- | --- | --- | --- |
| Data Set | CoV2_WT (control) | Cov2_P132H | Cov2_G15S | Cov2_K90R |
| HDX reaction details | 4C | 4C | 4C | 4C |
| HDX time course (sec) | 15, 60, 360,<br>3600 | 15, 60, 360,<br>3600 | 15, 60, 360,<br>3600 | 15, 60, 360,<br>3600 |
| # of peptides | 405 | 387 | 326 | 339 |
| Sequence coverage | 100.00% | 100.00% | 99.02% | 99.02% |
| Average peptide length / Redundancy | 17.24/22.82 | 17.29/21.86 | 16.33/17.39 | 16.65/18.44 |
| Replicates | 4 | 4 | 4 | 4 |
| Repeatability (avg. stddev of #D) | 0.1123 | 0.1200 | 0.1777 | 0.1412 |
| Significant differences in HDX (delta<br>HDX > X D - 95% CI) | n/a | 0.3123 D | 0.373 D | 0.3657 D |

## Data Availability

RCSB Protein Data Bank Accession code: The coordinates and structure factors have been deposited with the RCSB Protein Data Bank under accession codes: 7U29 (K90R), 7U28 (G15S) and 7TLL (P132H).

## Acknowledgements

We thank Mark Noe, Gretchen Dean, Annaliesa Anderson and Charlotte Allerton for critical reading of the manuscript and leadership.

